# Proximity-dependent biotinylation to elucidate the interactome of TNK2 non-receptor tyrosine kinase

**DOI:** 10.1101/2021.06.30.450607

**Authors:** Raiha Tahir, Anil K. Madugundu, Savita Udainiya, Jevon A. Cutler, Santosh Renuse, Li Wang, Nicole A. Pearson, Chris Mitchell, Nupam Mahajan, Akhilesh Pandey, Xinyan Wu

**Affiliations:** Biochemistry, Cellular and Molecular Biology Graduate Program, Johns Hopkins University School of Medicine, Baltimore, MD, USA; Department of Biological Chemistry, Johns Hopkins University School of Medicine, Baltimore, MD, USA; McKusick-Nathans Institute of Genetic Medicine, Johns Hopkins University School of Medicine, Baltimore, MD, USA; Institute of Bioinformatics, International Technology Park, Bangalore, 560066 India; Manipal Academy of Higher Education (MAHE), Manipal 576104, Karnataka, India; Pre-Doctoral Training Program in Human Genetics, McKusick-Nathans Institute of Genetic Medicine, Johns Hopkins University School of Medicine, Baltimore, MD, USA; Ginkgo Bioworks, Boston, MA, USA; Center for Molecular Medicine, National Institute of Mental Health and Neurosciences (NIMHANS), Hosur Road, Bangalore 560029, India; Departments of Pathology and Oncology, Johns Hopkins University School of Medicine, Baltimore, MD, USA; Department of Laboratory Medicine and Pathology, Mayo Clinic, Rochester, MN 55905, USA; Center for Individualized Medicine, Mayo Clinic, Rochester, MN 55905, USA; Molecular Pharmacology and Experimental Therapeutics, Mayo Clinic, Rochester, MN 55905, USA; Siteman Cancer Center, Washington University, St. Louis, MO, USA

## Abstract

Non-receptor tyrosine kinases represent an important class of signaling molecules which are involved in driving diverse cellular pathways. Although, the large majority have been well-studied in terms of their protein-binding partners, the interactomes of some of the key non-receptor tyrosine kinases such as TNK2 (also known as activated Cdc42-associated kinase 1 or ACK1) have not been systematically investigated. Aberrant expression and hyperphosphorylation of TNK2 has been implicated in a number of cancers. However, the exact proteins and cellular events that mediate phenotypic changes downstream of TNK2 are unclear. Biological systems that employ proximity-dependent biotinylation methods, such as BioID, are being increasingly used to map protein-protein interactions as they provide increased sensitivity in discovering interaction partners. In this study, we employed BioID coupled to the biotinylation site identification technology (BioSITe) method that we recently developed to perform molecular mapping of intracellular proteins associated with TNK2. We also employed stable isotope labeling with amino acids in cell culture (SILAC) to quantitatively explore the interactome of TNK2. By performing a controlled comparative analysis between full-length TNK2 and its truncated counterpart, we were not only able to confidently identify site-level biotinylation of previously well-established TNK2 binders and substrates such as NCK1, NCK2, CTTN, STAT3, but also discover several novel TNK2 interacting partners. We validated TNK2 interaction with one of the novel TNK2 interacting protein, clathrin interactor 1 (CLINT1), using immunoblot analysis. Overall, this work reveals the power of the BioSITe method coupled to BioID and highlights several molecules that warrant further exploration to assess their functional significance in TNK2-mediated signaling.

## INTRODUCTION

Signal transduction is an important component of intracellular events that drive biological processes. The relaying of signaling is largely driven by physical association of various classes of proteins, especially kinases. Early signaling events are mediated by the activity of tyrosine kinases, which include receptor tyrosine kinases (RTKs) that function at the cell surface as well as nonreceptor tyrosine kinases (NRTKs) that are predominately involved in cytoplasmic events. Activation of downstream signaling events by tyrosine kinases requires protein-protein interaction with intracellular binding partners as well as protein phosphorylation substrates. The functional relevance of signaling activities involving tyrosine kinases is evident from the phenotypic consequences of their dysregulation. Genomic events, such as amplification, rearrangement, or mutation, that led to aberrant activity of kinases have been associated with the development and progression of many subtypes of cancer as well as other diseases. Consequently, many therapeutic strategies for treatment of cancers involve the use of small molecule inhibitors or monoclonal antibodies that antagonize the activity of tyrosine kinases.

While most tyrosine kinases have been extensively studied to elucidate their cellular roles, including their protein interaction partners, some remain understudied despite evidence of their role in disease. One such kinase is the non-receptor tyrosine kinase TNK2 (also known as Activated CDC42-associated Kinase 1 or ACK1). Previous studies have revealed the role of the multiple protein domains of TNK2 in mediating its interaction with effectors and phosphorylation substrates, and the crucial role of these interactions in the regulation of TNK2 protein expression and kinase activity (1). The differential binding of TNK2 to various ubiquitin ligases, including NEDD4 (2), WWOX (3), and SIAH (4), through its PPXY motif and WW domain interacting region is involved in regulating the degradation of TNK2 protein. Various mechanisms and effectors are involved in regulating the activation status of TNK2. Through its MIG6 homology region (MHR), TNK2 is capable of binding a large variety of RTKs that are activated by multiple ligands, including EGF, GAS6, heregulin, IGF, and insulin. This growth factor-induced activation of TNK2 leads to various downstream events. TNK2 activation can induce the formation of homodimers, resulting in further activation of TNK2 via transphosphorylation (1). TNK2 can also be activated by SRC-mediated phosphorylation of the activation loop in the kinase domain (5). Furthermore, TNK2 is capable of auto-regulation through the inter-domain interactions of the kinase domain with its SH3, CDC/RAC-interactive (CRIB), proline-rich and MHR domains (6). TNK2 has been shown to interact with sequestosome 1 (p62/SQSTM1) through ubiquitin-association (UBA) domain for autophagy and binds to clathrin (CLTC) using its clathrin interacting region for endocytosis (7). Recent studies revealed that TNK2 can directly interact with and phosphorylate AKT1 and histone H4 and regulate their functions (8, 9). These interactions collectively place the role of TNK2 in pathways regulating the trafficking and recycling of receptor tyrosine kinases, endocytosis, cell adhesion, and other cellular processes (1).

Many studies have reported the involvement of aberrant TNK2 activity in driving various cancer subtypes. Activating mutation or overexpression of TNK2 are capable of driving the androgen-independent progression in castration-resistant prostate cancers (3, 10). Increased protein expression and hyperphosphorylation of TNK2 has been associated with poorer prognosis of patients with triple negative breast cancer (11, 12). Oncogenic mutations in TNK2 have been identified in leukemia (13). Other cancers where abnormal TNK2 activity was shown to promote disease include hepatocellular carcinoma (14), colorectal and gastric tumors (15), non-small-cell lung cancer (16), gliomas (17), and many other cancer subtypes. In addition to cancer, aberrant TNK2 activity has also been implicated in neurological problems such as Parkinson’s disease (18) and phenotypes involving abnormal dopamine transporter function (19). While individual signaling events, such as hyperphosphorylation of TNK2 substrates, have been used to explain the molecular mechanisms by which TNK2 promotes disease, a deeper understanding of these mechanisms requires knowledge of all potential molecules downstream of TNK2.

There are several methods to identify and characterize protein-protein interactions such as co-immunoprecipitation (co-IP) or pull-down assays. As a supplement to classic methods such as yeast two-hybrid (Y2H) or biophysical (X-ray, electron microscopy or NMR), recently developed BioID (proximity-dependent biotin identification) technology uses the promiscuous biotin ligase fused to the protein of interest (bait) in living cells, which can biotinylate the proximal interacting proteins followed by biotin capture and LC-MS/MS analysis. In our recent study, a modification made to the existing BioID workflow by quantitative BioSITe (<u>B</u>iotinylation <u>S</u>ite <u>I</u>dentification <u>T</u>echnology) of biotinylated peptides pulled down by anti-biotin antibody has largely expanded the scope of studying spatial and topological information of protein complexes (20–22). In the present study, we employed BioID coupled to BioSITe to map the interacting proteins of TNK2 in triple negative breast cancer cells. By using SILAC labeling-based quantitative proteomics strategy (23, 24) to differentially label full-length TNK2 and truncated version of TNK2, we were able to more clearly identify true binders of full-length TNK2 from false positive interactors and potential background contaminants. Collectively, our global study on the TNK2 interactome highlight a more specific approach to provide insights in oncogenic TNK2 signaling and identifying potential targets which may be used in combination therapy to apply multiple approaches of inhibiting TNK2 deregulated pathways or any compensatory/refractory mechanisms.

## METHODS AND MATERIALS

### Plasmids, cloning, antibodies, and reagents

Anti-biotin antibody (Bethyl Laboratories, #150-109A), streptavidin-HRP (Abcam, #ab7403), NCK1 Ab (CST, #2319S), HA-Tag Ab (Abcam, #ab9110), Clint1 Ab (Bethyl, #A301-926A), TNK2 Ab (Santa Cruz, sc-28336), Alexa Fluor 488 goat anti-mouse IgG (Invitrogen, A-11029), Alexa Fluor 568 goat anti-rabbit IgG (Invitrogen, A-11036), SlowFade™ gold antifade reagent with DAPI (Invitrogen, S36938). IgG Ab (CST, #3900S), Protein G Dynabeads (Thermo Fisher Scientific, #10003D), protein G beads (EMD Millipore, #16-266), biotin (Sigma Aldrich, #B4501), lipofectamine 2000 (Thermo Fisher Scientific, #11668019), sequencing grade trypsin (Promega, #V5113), and trypsin (Worthington Biochemical Corporation, #LS003741). SILAC amino acids (K8) and 13C6, 15N4-Arginine (R10) were from Cambridge Isotope Laboratories (Andover, MA), RPMI 1640 SILAC media deficient in L-lysine and L-arginine (Thermo Fisher Scientific, Waltham, MA). The plasmid pcDNA3.1 BioID-HA containing the mutant BirA-R118G was purchased through Addgene (Cambridge, MA, USA) (Plasmid #36047). The BirA*-HA cassette from above plasmid was used to generate pBABE BioID-HA plasmid. The full-length (FL) or N-terminal half (ΔC) of TNK2 sequence were cloned in-frame on the N-terminus of BioID-HA.

### Cell culture and generation of stable cell lines

HCC1395 triple negative breast cancer cells were transfected with pBABE-FL-BirA* and pBABE-ΔC-BirA* by overnight incubation in media containing plasmid and polybrene (1 ug/mL). Clones expressing protein of interest were derived using treatment and selection of transfected cells with puromycin.

### Co-Immunoprecipitation

FL and ΔC TNK2 HCC1395 cells were seeded in 150mm dishes and at confluence washed twice with PBS and harvested in co-IP buffer (20mM Tris HCL pH 7.5, 100mM NaCl, 1% NP40, 2.5 mM EDTA) and rotated gently for 1 hr at 4°C. The lysate was clarified by centrifugation at 15000 rpm for 15 mins and 2-3 mg of lysate was used per IP condition. 4-6 ug of HA-tag Ab (anti-rabbit Abcam) or IgG control Ab were added to corresponding cell lysates and incubated for 3 hours at 4°C followed by incubation with 50 ul of Dyna magnetic beads for 1 hr. Precipitated protein complexes were washed ice cold co-IP buffer for 4 times, eluted with 40 ul of 2X Laemmli buffer at 95°C for 5-10 mins and subjected to the Western blot.

### Quantitative SILAC proteomics for BioSITe

Two-state SILAC was performed for cells used in this study. Briefly, cells expressing mutant TNK2 ΔC-BirA* were labeled as “light” by culturing in media supplemented with light lysine (K0) and light arginine (R0). Cell expressing TNK2 FL-BirA* were labeled as “heavy” by culturing in media supplemented with ^13^C_6_, ^15^N_2_-Lysine (K8) and ^13^C_6_, ^15^N_4_-Arginine (R10). Cells expressing TNK2 FL-BirA* and TNK2 ΔC-BirA* were cultured overnight with 50 mM biotin. Cells were lysed in 8 M urea buffer (20 mM HEPES pH 8.0, 8 M urea, 1 mM sodium orthovanadate, 2.5 mM sodium pyrophosphate, 1 mM β-glycerophosphate, and 5 mM sodium fluoride), sonicated, and then cleared by centrifugation at 15,000 x g at 4 °C for 20 min. Protein concentration of lysates was determined by BCA Protein Assay. For each biological replicate, equal amounts of protein (10 mg) from each labeling condition was mixed and subjected to trypsin in-solution digestion and BioSITe analysis as described previously (21, 22, 25).

### Trypsin Digestion and Peptide Preparation

Briefly, lysate mixtures were reduced with 5 mM dithiothreitol and alkylated with 10 mM iodoacetamide. For in-solution tryptic digestion, the resulting protein extracts were diluted in 20 mM HEPES pH 8.0 to a final concentration lower than 2 M urea incubated with 1 mg/mL TPCK-treated trypsin on an orbital shaker at 25 °C overnight. Protein digests were acidified with 1% trifluoroacetic acid (TFA) to quench the digestion reaction and then subjected to centrifugation at 2000 xg at room temperature for 5 min. The resulting supernatants were desalted using SepPak C_18_ cartridge (Waters Corporation, Milford, MA). Eluted peptides were lyophilized to dryness prior to BioSITe analysis.

### BioSITe

Samples were processed using BioSITe method as previously described by Kim et al. (20) Briefly, peptide samples dissolved in BioSITe capture buffer (50 mM Tris, 150 mM NaCl, 0.5% Triton X-100) were incubated with anti-biotin antibody bound to protein-G beads for 2 hours at 4°C. Following incubation, beads were washed multiple times with PBS and then washed two times with BioSITe capture buffer, two times with 50 mM Tris and two times with ultrapure water. Biotinylated peptides were eluted four times using elution buffer (80% acetonitrile and 0.2% trifluoroacetic acid in water). The eluted sample was further cleaned up using C_18_ reversed-phase column and subject to LC-MS/MS analysis.

### LC-MS/MS analysis

Peptide samples were analyzed on an Orbitrap Fusion Lumos Tribrid Mass spectrometer coupled with the Easy-nLC 1200 nano-flow liquid chromatography system (Thermo Fisher Scientific). Peptides were reconstituted using 20 μl 0.1% formic acid and loaded on an Acclaim PepMap 100 Nano-Trap Column (100 μm x 2 cm, Thermo Fisher Scientific) packed with C_18_ particles (5 μm) at a flow rate of 4 μl per minute. Peptides were separated using a linear gradient of 7% to 30% solvent B (0.1% formic acid in 95% acetonitrile) at a flow rate of 300-nl/min over 95 min on an EASY-Spray column (50 cm x 75 μm ID, Thermo Fisher Scientific) packed with 2 μm C_18_ particles, which was fitted with an EASY-Spray ion source that was operated at a voltage of 2.3 kV Mass spectrometry analysis was performed in a data-dependent manner with a full scan in the mass-to-charge ratio (m/z) range of 300-18,000 in the “Top Speed” setting, three seconds per cycle. MS scans for precursor ion detection were measured at a resolution of 120,000 at an m/z of 200. MS/MS scans for peptide fragmentation ion detection were acquired by fragmenting precursor ions using the higher-energy collisional dissociation (HCD) method and detected at a mass resolution of 30,000, at an m/z of 200. Automatic gain control for MS was set to one million ions and for MS/MS was set to 0.05 million ions. A maximum ion injection time was set to 50 ms for MS and 100 ms for MS/MS. MS was acquired in profile mode and MS/MS was acquired in centroid mode. Higher-energy collisional dissociation was set to 32 for MS/MS. Dynamic exclusion was set to 35 seconds, and singly charged ions were rejected. Internal calibration was carried out using the lock mass option (m/z 445.1200025) from ambient air.

### Bioinformatics data analysis

Proteome Discoverer (v 2.2; Thermo Scientific) suite was used for identification. Raw files derived from 3 replicate LC-MS/MS runs, and were searched together. Spectrum selector was used to import spectrum from raw file. During MS/MS preprocessing, the top 10 peaks in each window of 100 m/z were selected for database search. The tandem mass spectrometry data were then searched using SEQUEST algorithm against protein databases (For BioID experiments; Human RefSeq database (v73 containing 73,198 entries) with the addition of FASTA file entries for TNK2 FL-BirA* and TNK2 ΔC-BirA*. The search parameters for identification of biotinylated peptides were as follows: a) trypsin as a proteolytic enzyme (with up to three missed cleavages); b) minimum peptide length was set to 6 amino acids. c) peptide mass error tolerance of 10 ppm; d) fragment mass error tolerance of 0.02 Da; and e) carbamidomethylation of cysteine (+57.02 Da) as a fixed modification and f) oxidation of methionine (+15.99 Da), ^13^C_6_, ^15^N_2_-lysine (K8),^13^C_6_, ^15^N_4_-arginine (R10), biotinylation of lysine (+226.07 Da), biotinylation of heavy lysine (+234.09) as variable modifications. Peptides were filtered at a 1% false-discovery rate (FDR) at the PSM level using percolator node.

Identified protein and peptide spectral match (PSM) level data were exported as tabular files from Proteome Discoverer 2.2. We used PyQuant (26) for obtaining the relative quantification of biotinylated peptides with SILAC light or heavy amino acids. To derive the precursor ion abundance values for the isotopic counterparts for each biotinylated peptide identified from our database search, peak area calculated by PyQuant was used. Briefly, PyQuant scans for the light and heavy peptide (K8R10) peaks in the full MS1 scans and calculates the area under the curve to derive the peak area over the extracted ion chromatogram (XIC). Precursor ion abundances thus computed were used for relative abundance estimates of biotinylated peptides. Missing abundance values in any LC-MS/MS replicate experiment were replaced with the minimum value of the peptide peak area obtained from other replicates. We used an in-house Python script to compile the peptide level site information mapped to UniProt or RefSeq protein sequences. The summary count on number of biotinylated sites, supported peptides and PSMs and the relative fold-change information are then calculated at peptide and protein levels.

### Immunofluorescence staining

Briefly, cells were cultured in 8-well chamber slides. After fixing with 4% formalin in PBS, cells were incubated overnight at 4°C with anti-TNK2 mouse monoclonal antibody (1:100) and anti-CLINT1 rabbit polyclonal antibody (1:100), or with anti-TNK2 mouse monoclonal antibody (1:100) and anti-HA rabbit monoclonal antibody (1:200). Next day, cells were washed PBS for three times and secondary antibodies (Alexa Fluor 488 goat anti-mouse IgG, Alexa Fluor 568 goat anti-rabbit IgG) were added and incubated at 37°C for 1 hour with 1:500 dilution. After PBS wash, the slides were sealed with SlowFade™ gold antifade reagent with DAPI. Images were taken by Zeiss LSM 980 confocal microscope with Airyscan 2.

### Data availability

All mass spectrometry datasets acquired for this study were deposited to ProteomeXchange (http://proteomecentral.proteomexchange.org) and is available via the PRIDE database with the accession number PXD010474. Reviewers can access the dataset by using as ID and “Z64u9xFz” as password.

## RESULTS AND DISCUSSION

The dynamic assembly and disassembly of protein complexes is a fundamental principal that drives biological processes. Here, we describe quantitative analysis of protein biotinylation with a site-level resolution to identify TNK2 proximal proteins in the context of a breast cancer cell line.

### Application of BioSITe for the discovery of TNK2 interacting proteins

TNK2 is a non-receptor tyrosine kinase with a multidomain structure. TNK2 is composed of at least eight distinct domains: the sterile α motif (SAM), kinase domain, Src homology domain 3 (SH3) domain, GTPase-binding domain, clathrin-interacting region, WW domain-interacting region, an MIG6 homology region (MHR), also known as epidermal growth factor receptor (EGFR)-binding domain (EBD) and an ubiquitin association (UBA) domain (Figure 1A). The SAM domain and kinase domain reside in the N-terminal moiety of TNK2. The C-terminal part of TNK2 consists of multiple domains mediating TNK2 interaction with other proteins, including clathrin-interacting region, WW domain-interacting region, MHR, and UBA domain To study the interactome of TNK2, we took advantage of the BioID system to identify interacting proteins of TNK2. For this, we generated a full-length (FL) TNK2 construct fused with BirA* at C-terminal of TNK2 **(Figure 1A)**. We also generated a TNK2 C-terminal deletion BirA* fusion construct (ΔC) consisting of the N-terminal moiety of TNK2 (1 - 444 AA), with the deletion of the C terminal moiety of TNK2 (ΔC) that was lacking the clathrin-interacting region, the Proline-rich domain, MHR, EBD and UBA domains. This truncated construct also served as a control to allow us to more clearly differentiate the interacting partners of the functional full-length TNK2 protein. The full length TNK2 (FL) and truncated TNK2 (ΔC) constructs were respectively transfected into HEK293T cells for packaging viral particles that were used to infect HCC1395 cells and stable cell lines were generated by puromycin selection. Relative expression levels of BirA*-tagged TNK2 FL, and ΔC were analyzed by immuno-blot with both anti-HA tag and anti-TNK2 antibodies, showing the different molecular weights of TNK2 protein variants. Western blot analysis using an HA tag and TNK2 specific antibody confirmed the expression of both constructs at their expected molecular weights **(Figure 1B)**. We also performed immunofluorescent (IF) staining for HCC1395 cells expressing ΔC-TNK2-BirA-HA. Anti-HA tag antibody was used to detect exogeneous △C-TNK2-BirA-HA and anti-TNK2 mAB (SCBT: A-11) that recognizes TNK2 C-terminal moiety, was used to detect the expression endogenous TNK2. Our result confirmed that △C-TNK2 is largely colocalized with endogenous full-length TNK2 (Figure S1B). To confirm an increase in global intracellular biotinylation in cells with stable expression of TNK2 BirA* fusion constructs, we performed western blot analysis using anti-biotin antibody on whole cell lysates from stable cells cultured in growth medium with or without exogenous biotin overnight. As expected, relative to parental cells and cells not treated with biotin, our BirA* cells incubated with biotin showed an increased global biotinylation profile, which included visibly increased biotinylation of the TNK2-BirA* fusion proteins themselves **(Figure 1C)**. To systematically map the interacting proteins specific to the canonical full-length TNK2 protein, we opted for a quantitative approach that employs stable isotope labeling amino acids in cell culture (SILAC) (27). The stable cell lines with FL TNK2 or ΔC TNK2 were labeled by growing in heavy (K8, R10) and light (K0, R0) amino acids supplemented SILAC media, respectively, and treated overnight with exogenous biotin **(Figure 1D)**. The cells were harvested, lysed, mixed and then digested with trypsin. To identify proteins biotinylated downstream of full-length or truncated TNK2 protein, we employed a method recently developed in our lab that allows the site-specific analysis of biotinylation on peptides via mass spectrometry analysis (18). Using whole cell lysates derived from SILAC-labeled breast cancer cells (HCC1395) expressing either ΔC mutant (Light) or wild-type (Heavy) TNK2-BirA*, we performed immunoaffinity-based enrichment of biotinylated peptides using an anti-biotin antibody.

**Figure 1.**
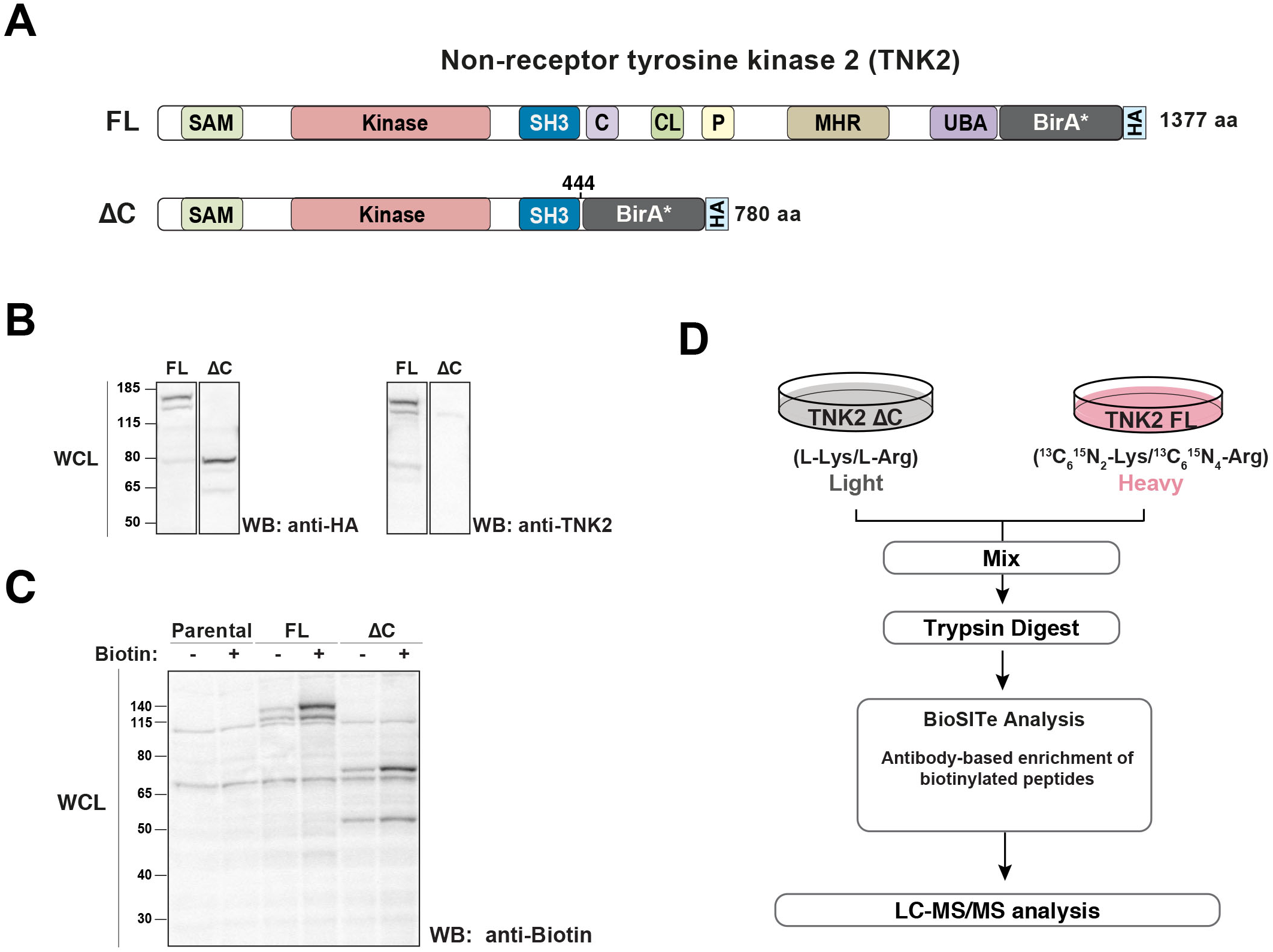
Overview of SILAC-BioSITe quantitative workflow for mapping TNK2 interactome in breast cancer cells. (A) Schematic depicting the protein domains within recombinant TNK2 full length (top) and ΔC mutant (bottom) constructs used in this study. A biotin ligase (BirA) and HA tag were cloned in-frame at the C-terminus of both variants as indicated. (B) Western blot analysis of whole cell lysates from breast cancer cells (HCC1395) to confirm expression of TNK2 FL-BirA* or ΔC-BirA*constructs. Analysis was performed using antibodies against TNK2 and HA-tag. (C) Western blot analysis of global biotinylation using whole cell lysates from parental cells and cells expressing TNK2 FL-BirA* or ΔC-BirA*. Analysis was performed using a biotin-specific antibody. (D) Experimental workflow for differential interactome analysis of SILAC-labeled HCC1395 cells expressing full-length (Heavy) or ΔC mutant (Light) TNK2-BirA* fusion constructs were incubated overnight with media containing biotin. Equal amounts of cell lysates from each condition were mixed and digested into peptides. Biotinylated peptides were enriched using BioSITe and analyzed by LC-MS/MS.

### Mass spectrometry and data analysis to identify and quantify biotinylated peptides and proteins

As part of our strategy of applying proximity-based biotinylation for mapping the intracellular interacting proteins of TNK2, we acquired quantitative mass spectrometry data from the analysis of enriched fractions of biotinylated peptides derived from three biological replicates. LC-MS/MS analysis using an Orbitrap Fusion Lumos generated raw mass spectrometry data that were subjected to spectral matching via a combined database search. Our experimental design called for a search scheme that could identify SILAC labeled or unlabeled peptides modified with biotin at one or more/all lysine residues. To consider all such peptides, we configured our database search parameters to consider the presence of multiple variable modifications, including SILAC modification of lysine and arginine (Lys8, Arg10), biotinylation of light (Lys-Biotin) as well as SILAC-modified lysine (Lys8+Biotin) **(Figure 2A)**. We also allowed for 3 missed cleavages to account for the lack of cleavage by trypsin at C-terminus of lysine residues with biotinylated sidechains. Database spectral matching with the above parameters led to the identification of 751 unique biotinylated peptides (Supplemental Table S2) corresponding to 356 biotinylated proteins (Supplemental Table S3). When possible, MS/MS spectra were manually examined to check for the presence of signature fragment ions, such as those resulting from biotinylated heavy (m/z = 316.17) and light (m/z = 310.15) lysine residues(21), and to confirm the presence and quality of fragments covering the annotated biotinylation site.

**Figure 2.**
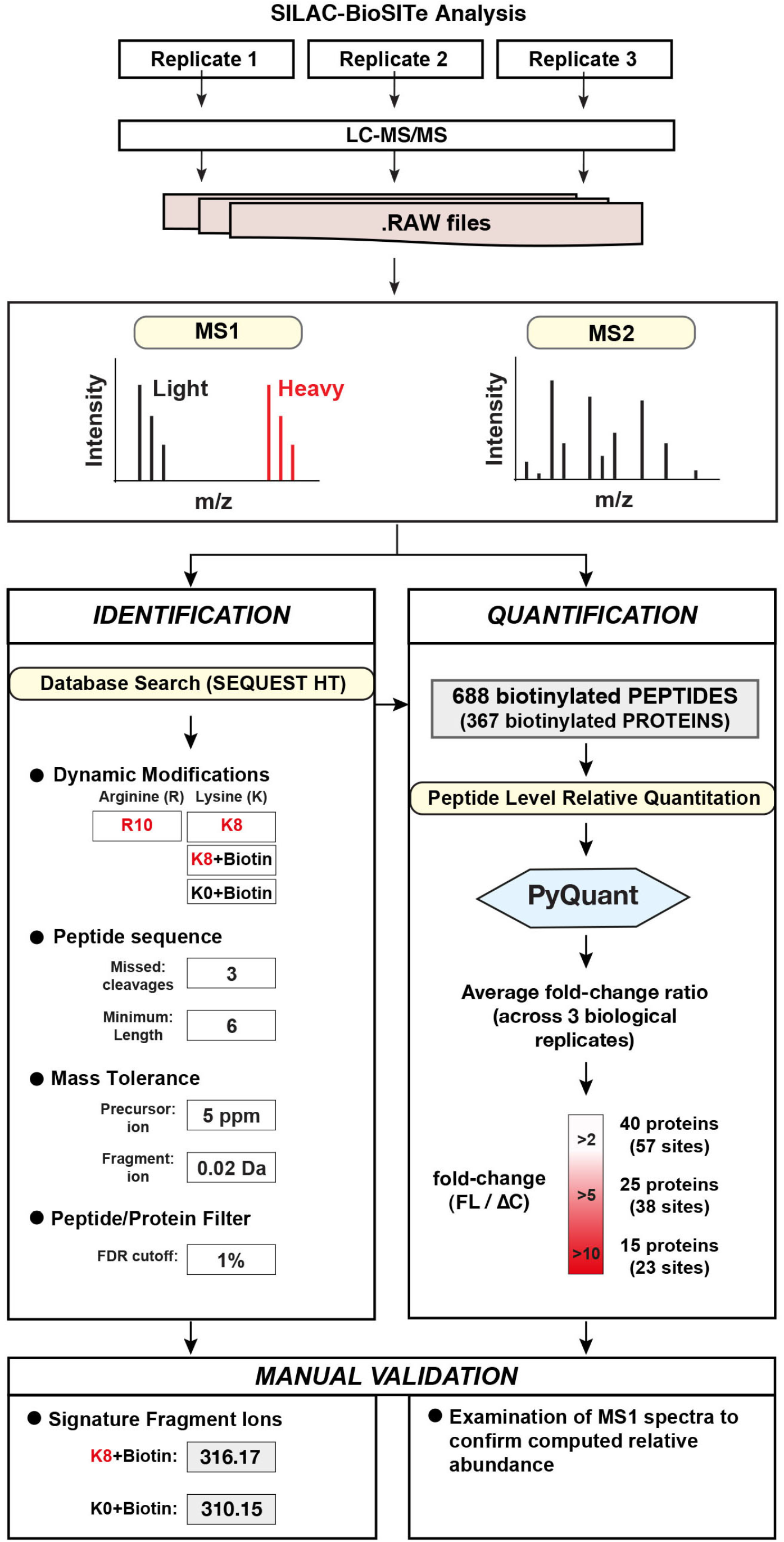
Data analysis pipeline to identify and quantify biotinylated peptides. The overall computational pipeline used for processing raw files to identify and quantify biotinylated peptides is shown. Raw files were processed to generate a list of biotinylated peptide identifications. The parameters used for peptide identification using SEQUEST included the various dynamic modification and mass tolerance thresholds as indicated. Quantification was performed using PyQuant (28). The number of peptides (and proteins) identified and their corresponding foldchange values are indicated.

The list of biotinylated peptides identified from the combined use of SILAC-labeling with BioSITe called for a more versatile quantitation pipeline. As depicted in **Figure 2A**, the biotinylated peptides in our study could occur in multiple forms and could be isotopically labeled (K8), biotinylated, or both. With the added consideration of mis-cleaved peptides, these parameters collectively lead to a substantial increase of the complexity for isotope precursor ion identification and quantification, resulting in failure to quantify certain peptides with multiple dynamic modifications.. Using a quantification workflow that integrated PyQuant, a quantification algorithm previously developed by our group (28), we derived MS1 level quantification for the list of identified biotinylated species and their corresponding isotopic counterparts **(Figure 2A)**. This enabled us to get quantitation for 749 (out of 751) identified biotinylated peptides. Representative MS2 spectra along with the corresponding MS1 spectrum used for deriving relative quantification are shown for biotinylated peptides from NCK interactor 1 (NCK1) **(Figure 3A)**, Cortactin (CTTN) **(Figure 3B)**, Clathrin interactor 1 (CLINT1) (Figure 4.3C) and TELO2-interactor 1 (TTI1) protein **(Figure 3D)**.

**Figure 3.**
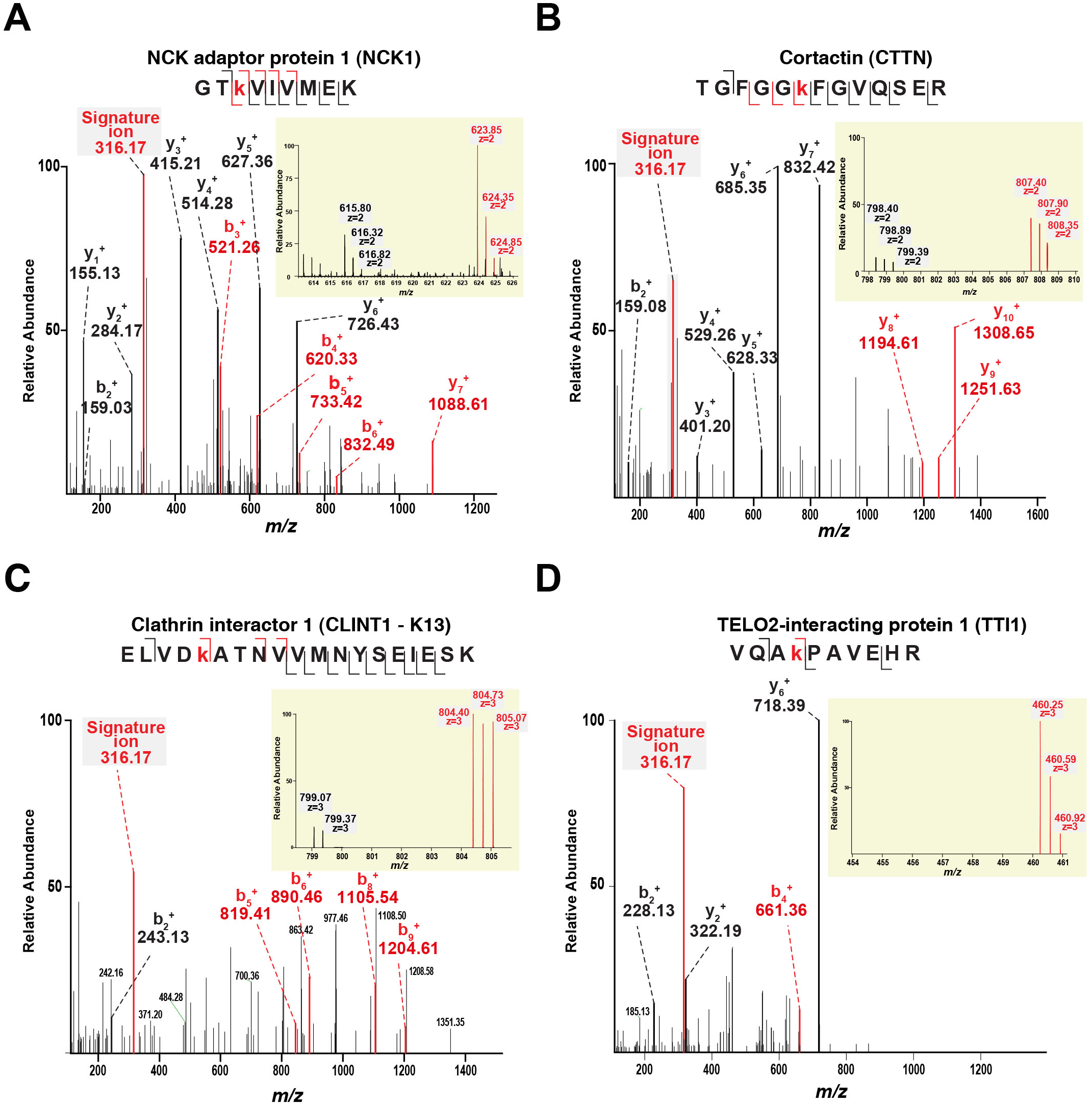
Representative MS/MS and MS spectra used for identification and relative quantitation of biotinylated peptides derived from cells expressing TNK2-BirA* constructs. (A) MS/MS fragmentation of a biotinylated peptide mapping to NCK adaptor protein 1 (NCK1), a known binding partner of TNK2. Representative MS1 spectrum used for deriving relation quantification between full-length and ΔC mutant cells is shown as an inset. Fragments ion that confirm the annotated site of biotinylation are shown in red. Signature fragment ions resulting from biotinylated lysine moiety are indicated in green. (B) Representative MS/MS and MS1 spectra of biotinylated peptide mapping to K272 in cortactin (CTTN), a known interactor and phosphorylation substrate of TNK2. (C) Representative MS2 and MS spectra of biotinylated peptide mapping to selected novel TNK2 interacting proteins such as (C) clathrin interactor 1(CLINT1) and (D) TELO2-interacting protein 1 (TTI1).

**Figure 4.**
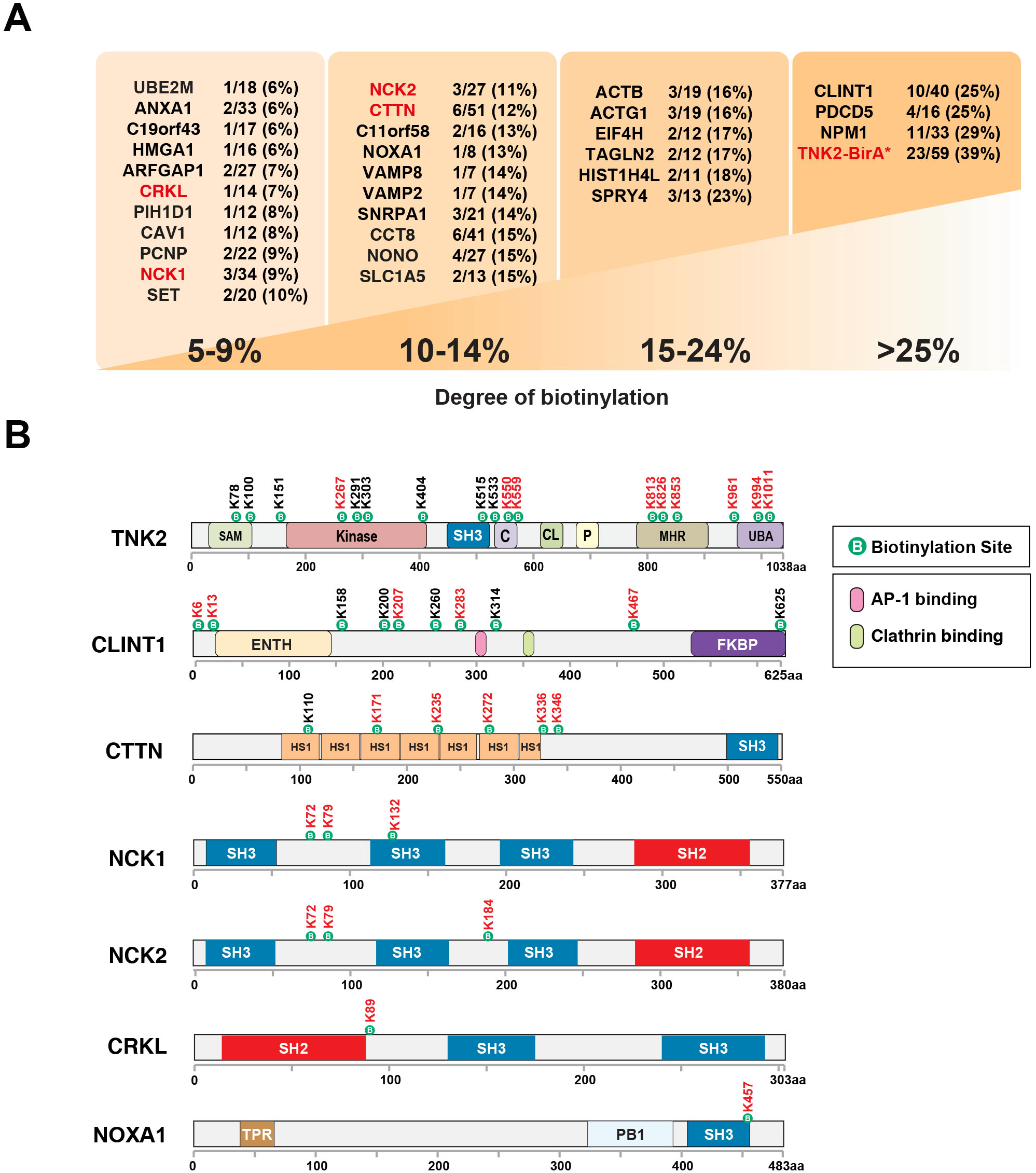
Mapping of TNK2 interactome using biotinylation sites. (A) Biotinylated proteins identified by BioSITe are shown grouped by the degree of biotinylation, defined by the number of biotinylated lysines out of the total number of lysines in each protein. Known binding partners of TNK2 are shown in red. (B) Domain schematics for select biotinylated proteins identified in this study. The protein domains and the relative localization of the biotinylation sites mapped to the protein are shown. The full names of the protein domains are described in the legend. Sites showing a 2-fold or greater change in abundance ratio between TNK2 FL-BirA* and TNK2 ΔC-BirA* are labeled in red.

### Identification of TNK2 proximal interactome using BioSITe

Our group has previously shown that the degree of protein biotinylation can be used as a potential metric to help prioritize confident interacting proteins from potential contaminants and false positives (22, 25). Therefore, we grouped the biotinylated proteins in our data based on their degree of biotinylation **(Figure 4A)**. As expected, this resulted in the sorting of TNK2 bait protein as the protein showing the highest degree of biotinylation. TNK2 is known to bind to itself and form homodimers via its N-terminal SAM domain (29). Therefore, in addition to autologous biotinylation, identification of biotinylated peptides mapping to TNK2 may in part be a result of TNK2 self-dimerization. In search for true interaction partners of TNK2, we next examined the set of biotinylated sites and proteins that showed a higher abundance (>= 2 fold) in cells expressing full-length TNK2 (FL-TNK2) relative to cells expressing a truncated version of the protein (ΔC-TNK2). Considering that ΔC-TNK2 with the intact N-terminal domains of TNK2, ΔC-TNK2 would be able to bind to endogenous TNK2 and form dimmers in HCC1395 cells. It is possible that the BirA domain fused to ΔC-TNK2 could transfer biotin to the proteins associated with endogenous full-length TNK2. In this case, the calculated ratio would be slightly underestimated, resulting in missing identification of some of the TNK2 associated proteins. However, those biotinylated proteins with higher biotinylation (>=2 fold) in FL-TNK2 cells than in ΔC-TNK2 cells are likely the bona fide TNK2 associated proteins. Our analysis led to the identification of 95 biotinylation sites mapping to 48 biotinylated proteins that displayed a ≥2-fold higher abundance in cells expressing full-length relative to truncated protein (referred to from here on as full length specific TNK2 binders). Among these were biotinylation sites mapping to several experimentally established TNK2 binders, including NCK adaptor protein 1 (NCK1), NCK adaptor protein 2 (NCK2) (5, 30), cortactin (CTTN) (31), clathrin (CLTC) (7), and Cyclin G associated kinase (GAK). Some of our biotinylated candidates overlapped with those identified in previous co-immunoprecipitation experiments using recombinant TNK2, including STAT3 (K707) and CRKL (K89). Analysis in the previous study also showed that TNK2 was involved in phosphorylation of STAT3 at Y705, and that this process is partly regulated by HSP90 (32). In line with these reports, we also observed increased biotinylation of several heat shock protein complex components in the cells expressing full-length TNK2-BirA*, including HSP8 (K507, K512), HSP90AB1 (K607), and HSPA1A (K507). Furthermore, while CRKL was identified in the above study, we also found biotinylated peptides mapping to CRK, which is not a known TNK2 binder. Overall, these findings indicate that the relative abundance ratio for biotinylation sites/peptides between full-length and truncated constructs can serve as a powerful differentiator between true binders of the biotin ligase fused bait protein and background or false positives. In this study, this strategy allowed us to identify true binders of TNK2 protein by relative comparison of protein biotinylation between cells expressing full-length TNK2 and truncated TNK2.

In addition to identifying sites of biotinylation on several proteins that overlap with published interacting proteins and substrates of TNK2, our BioSITe data also included many biotinylated proteins that have not been previously reported to be associated with TNK2 and could be potentially novel TNK2 interacting proteins. These interaction candidates included clathrin interactor 1 (CLINT1), TELO2 interacting protein 1 (TTI1), NADPH oxidase activator 1 (NOXA1), RuvB like AAA ATPase 1 (RUVBL1), and GRB10 interacting GYF protein 2 (GIGYF2). We noted that several of the novel interaction candidates for TNK2 are proteins that bind known interacting proteins of TNK2, as determined by previously reported studies. For instance, comparison of 101 known CLINT1 interacting proteins with known TNK2 interacting proteins indicates common binding to several well-characterized TNK2 interacting proteins, included CLTC, GAK, NCK1, GTSE1, and NTRK1. Similarly, in the database derived interactome for TTI1 (n = 47), an important member of the mTOR protein complexes (33), binds EGFR and NTRK1.

TNK2 is known to bind SH3 domains through its proline rich domain (34–36). To test for an enriched presence of SH3 and potentially other protein domains that mediated molecular association with proteins, we performed analyses to determine whether proteins showing fulllength specific biotinylation were enriched in certain protein domains, and also whether the site of biotinylation in these putative TNK2 binding partners were preferentially localized to specific protein domains and regions. A more in-depth examination of biotinylation site localization in the context of the annotated domains in individual proteins revealed that some sites mapped onto or adjacent to known protein domains and motifs, namely SH3 domains (CRK, CTTN, DNMBP, NCK1, NCK2, NOXA1, STAM2, TNK2, UBASH3B) and the ENTH domain which is involved primarily in mediating binding to membrane lipids (CLINT1, HGS, HIP1, STAM2, TTI1). A few selected biotinylated proteins and the location of biotinylation sites with their linear protein structure is depicted in **Figure 4B**. Furthermore, we observed that while several proteins in our data were found biotinylated at multiple sites, the relative abundance of the biotinylated peptides mapping to regions adjacent to these protein domains were often more abundant in cells expressing full-length TNK2 where the proline-rich domain is conserved.

### TNK2 proximally biotinylated proteins are involved in vesicle-mediated endocytosis

A collective survey of previous studies available on TNK2 indicates its association with pathways involved in recycling of surface receptors, especially receptor tyrosine kinases such as Epidermal growth factor receptor (EGFR), Insulin receptor, and AXL receptor tyrosine kinase (37–39). However, the specific proteins and protein complexes engaged by TNK2 to drive these processes have not been properly surveyed previously. Thus, we performed gene ontology (GO) enrichment analysis to determine whether proteins interacting with full length TNK2 were known to be involved with these and other novel biological pathways and processes **(Figure 5A)**. Survey of biological processes and KEGG pathways revealed an enrichment of proteins mapping to pathways associated with either endocytosis (CLINT1, CLTC, CTTN, GAK, HIP1, HSP90AA1, HSPA8), receptor tyrosine kinase signaling (GIGYF2, HGS, HSP90AA1, STAM2, STAT3), or both (CRK, EPS15L1, NCK1, TNK2) **(Figure 5A)**. Next, we sought to investigate TNK2 protein interaction network using STRING analysis. This analysis revealed that a large majority of the proteins were highly connected to other proteins identified in this study as TNK2 interacting proteins. Only 8 proteins did not have any previously described interaction with any of the other proteins in the list of TNK2 interacting proteins. Significantly, nine of the proteins identified were known interacting proteins of TNK2 **(Figure 5B)**. Four major functional clusters or interaction modules were identified within the larger network potentially indicating proximal protein complexes. The top scoring cluster included three known TNK2 interacting proteins - CTTN, CLTC, and GAK – in addition to several novel highly interconnected interacting proteins - AGFG1, CLINT1, HIP1, HSP8, OCRL, and VAMP2 **(Figure 5C)**. Interestingly, this cluster accounted for the majority of proteins that were found in GO enrichment analysis to be associated with endocytosis and related processes indicating a potential protein complex that may be involved in carrying out TNK2-mediated endocytosis.

**Figure 5.**
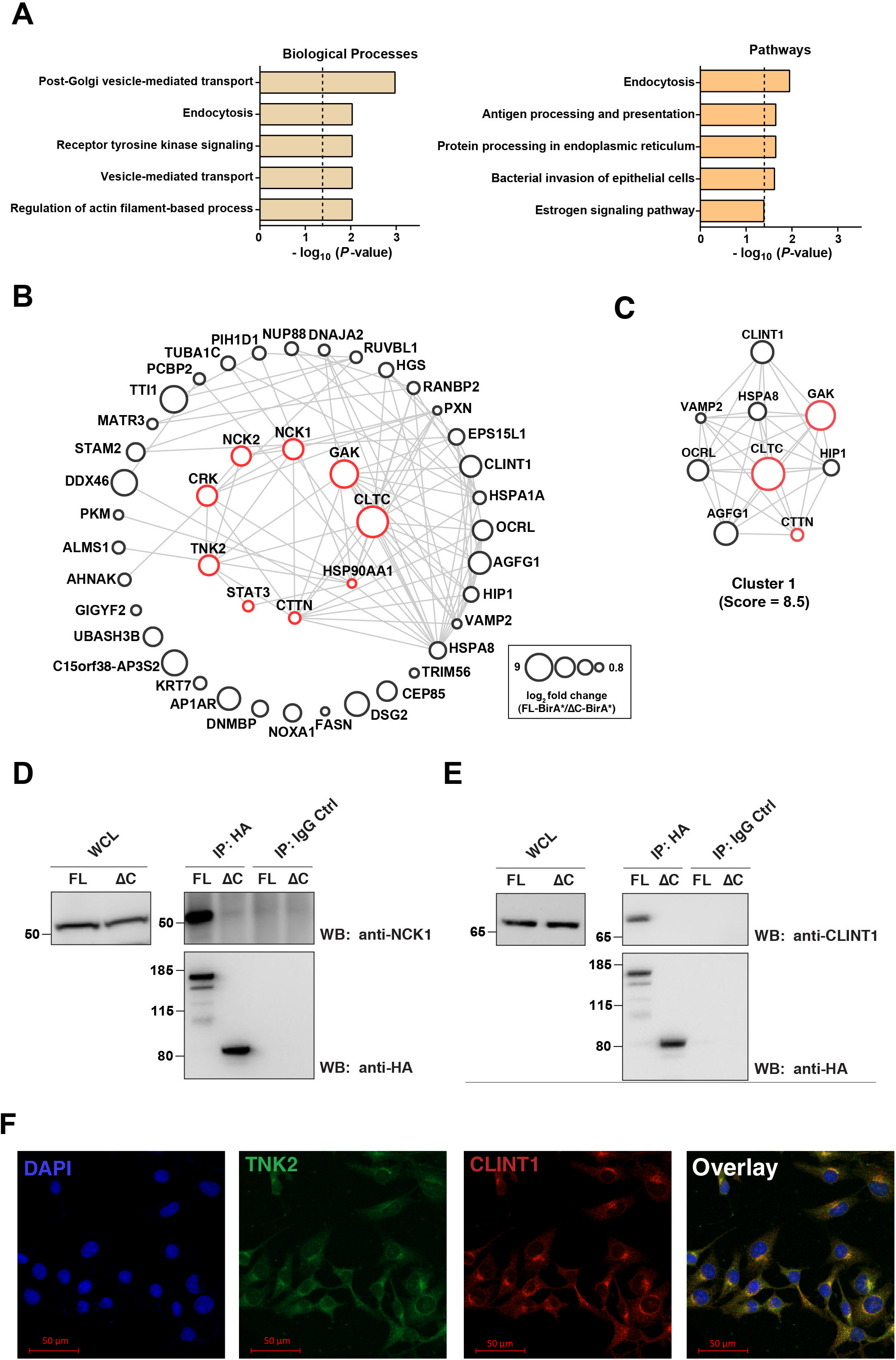
Pathway analysis and validation of TNK interacting proteins. (A) Results from Gene Ontology (GO) enrichment analysis of biotinylated proteins that displayed a ≥2-fold increase in abundance in cells expressing full-length TNK2 relative to its truncated ΔC version (FL/ΔC). Top five biological processes and KEGG pathways found to be significantly enriched (p < 0.05) are shown. Dashed line denotes p-value threshold for 0.05. (B) Depiction of STRING analysis of biotinylated proteins that displayed a ≥2-fold increase in abundance (FL/ΔC) performed using Cytoscape. The size of the circles representing the nodes is proportional to the protein level foldchange (FL/ΔC) as depicted. Red circles represent known interacting proteins of TNK2, or its downstream substrates previously demonstrated to be phosphorylated by TNK2. (C) A detailed view of the top scoring cluster from the protein interaction network shown in panel B. (D) The left panel shows Western blot analysis of whole cell lysates (WCL) expressing TNK2 FL-BirA* (FL) or ΔC-BirA* (ΔC). The panels on the right show a co-immunoprecipitation experiment using antibodies against HA-tag (HA) or a matched isotype control (IgG control) for immunoprecipitation (IP). The top right panel shows immunoblotting with anti-NCK1 antibody while the lower right panel shows immunoblotting with anti-HA-tag antibody. (E) Western blot analysis of control mouse IgG and anti-HA tag IP from cells expressing TNK2 FL-BirA* or ΔC-BirA*. Western blot analysis was performed using antibodies against Clathrin interactor 1 (CLINT1) and HA-tag. Analysis of whole cell lysates with anti-CLINT1 was performed in parallel.

In order to validate the findings of BioID-based interactome study using another approach, we performed coimmunoprecipitation assays to confirm the interactions between TNK2 and NCK1, a well characterized TNK2 interactor and Clathrin interactor 1 (CLINT1), a novel TNK2 interactor discovered by this study. NCK1 was > 50-fold more biotinylated in cells expressing full-length (FL) TNK2-BirA* fusion protein comparing to cells expressing truncated TNK2 (ΔC)-BirA fusion protein. Our coimmunoprecipitation experiments demonstrated that enrichment using anti-HA tag antibody could specifically coimmunoprecipitate NCK1 with HA-tagged FL TNK2-BirA*, but not with truncated HA-tagged TNK2 (ΔC)-BirA* fusion protein (Figure 5D). CLINT1 was discovered as a novel TNK2 interactor and was 12 time more biotinylated by FL-TNK2-BirA* than by TNK2 (ΔC)-BirA*). As shown in Figure 4B, we identified 10 biotinylated lysine sites in CLINT1, and all of them are more (>2) biotinylated by FL-TNK2-BirA* than by TNK2 (ΔC)-BirA*). These sites are scattered across the entire protein of CLINT1. In our validation study, CLINT1 was observed only in eluates obtained from anti-HA tag IP from lysates of cells expressing HA tagged FL-TNK2-BirA*, suggesting this interaction required the full-length TNK2 protein. We also performed immunofluorescence (IF) staining to examine the subcellular localization patterns of endogenous TNK2 and CLINT1 expressed in MDA-MB-231 cells. We observed overlap of fluorescent signals from TNK2 and CLINT1 concentrated at one corner outside of nucleus, indicating the colocalization of TNK2 and CLINT1 (Figure 5F). Therefore, our analysis demonstrates that a differential interactome study using a SILAC-BioSITe approach can enable the detection and identification of true novel interacting proteins of a protein of interest.

## CONCLUSIONS

Large-scale interaction studies employing high-throughput platforms have become increasing popular for obtaining “comprehensive” protein-protein interactome maps of protein families (40) and even entire human interactomes (41, 42). However, there is a need for appropriate controls or complementary methods that help pinpoint true interacting proteins among the large list of candidates rendered from one method alone. BioID and other proximity-based labeling strategies can serve as good complementary methods to assist in the prioritization and sorting of high confidence interacting proteins. In fact, large-scale studies based on proximity dependent biotinylation are already becoming popular for mapping interactomes for protein families, including G protein coupled receptors (43, 44), protein phosphatases (45), nuclear transport receptors (46), as well as for individual proteins like paxillin (47), fibroblast growth factor receptors (FGFRs) (48, 49), transcription factor SOX2 (50), dynein (51), and many others. However, these methods are not being applied for studying molecules whose protein interactomes have not yet been comprehensively characterized. In the present work, we described the interactome map of one of the few understudied but important kinases: non-receptor tyrosine kinase 2 (TNK2). TNK2 has an established role in driving various cancers, and we also recently identified it as one of the proteins that exhibits increased tyrosine hyperphosphorylation downstream of two different oncogenic KRAS mutations (52). We show the combined use of BioID with BioSITe, which allowed us to obtain site-level biotinylation evidence for putative TNK2 binders. Furthermore, using a recently developed proteomic quantification algorithm, PyQuant, we described a novel workflow that allows quantitative proteomics experiment employing SILAC with BioID-BioSITe to compare the relative abundance of site specific biotinylation modified by the bait protein or its control. This allowed us to identify known and novel TNK2 interacting proteins that warrant further experiments to assess their functional significance in TNK2-mediated signaling. We believe that the strategy used in this work is broadly applicable for the field of protein-protein interactions and should be applied to map the interactomes of other proteins of interest.

## ACKNOWLEDGEMENTS

This work was supported by the Wellcome Trust/DBT India Alliance Margdarshi Fellowship [IA/M/15/1/502023], NIH grant (R01CA184165) and a DOD breast cancer research breakthrough award (W81XWH-15-1-0311) to A.P. This work was also supported by the Mayo Clinic Breast Specialized Program of Research Excellence (SPORE) (P50CA116201) Developmental Research Program Award and a DOD breast cancer research breakthrough award (BC181309) to X.W.

## AUTHOR CONTRIBUTIONS

R.T., X.W., A.P. designed research; R.T., J.A.C., S.R., S.U., W.L., N.A.P. performed experiments; R.T., A.K.M., X.W. analyzed data; R.T., S.U., X.W., A.P. wrote the paper.

## SUPPLEMENTAL TABLES

Supplemental Table S1 – A complete list of PSMs with SILAC quantification calculated by Pyquant

Supplemental Table S2 – A complete list of biotinylated peptides identified in this study.

Supplemental Table S3 – A complete list of biotinylated proteins identified in this study

**Supplementary Figure 1.**
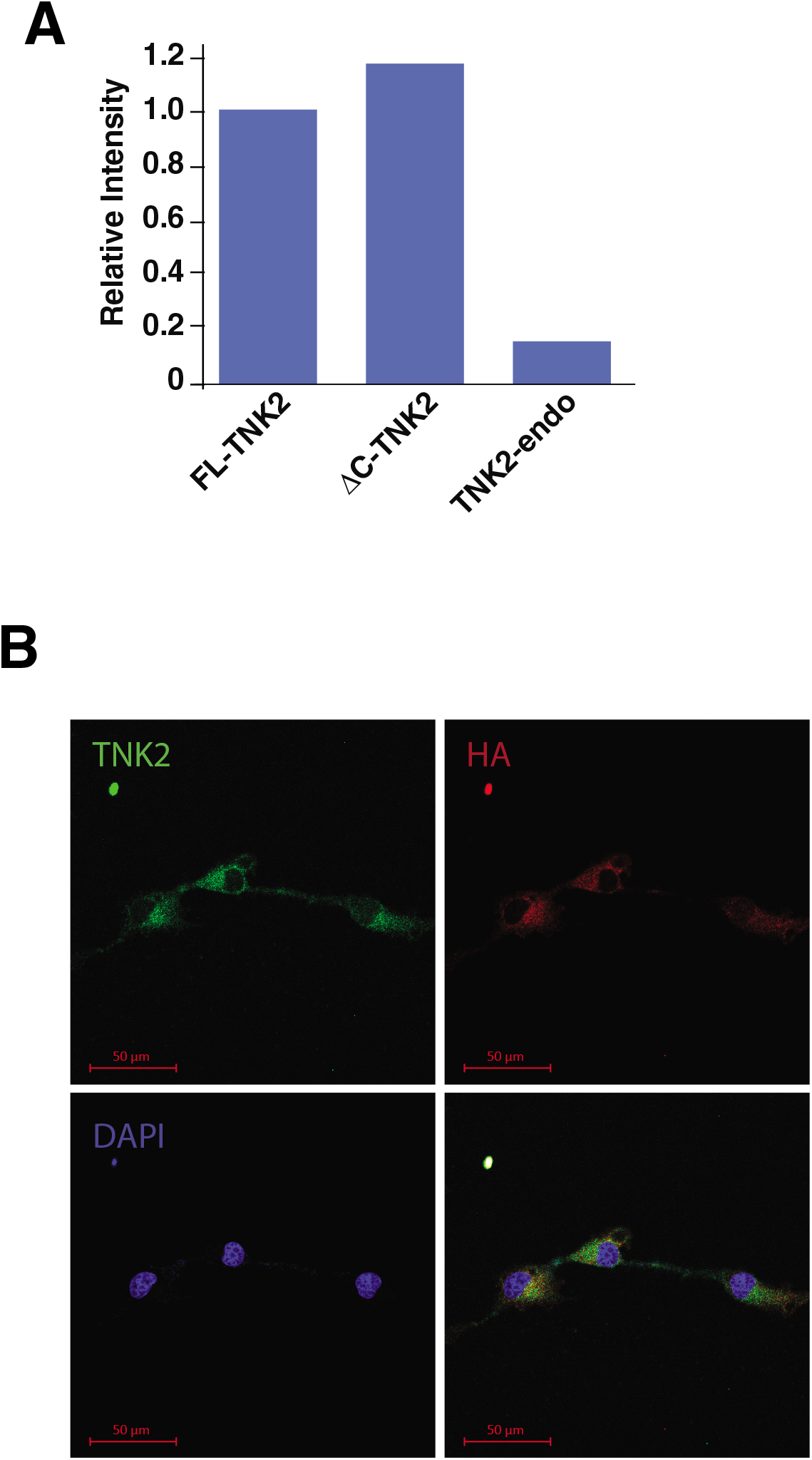

